# *miRNA-34c* suppresses osteosarcoma progression *in vivo* by targeting Notch and E2F

**DOI:** 10.1101/2021.05.24.445452

**Authors:** Yangjin Bae, Huan-Chang Zeng, Yi-Ting Chen, Shamika Ketkar, Elda Munivez, Zhiyin Yu, Francis H. Gannon, Brendan H. Lee

## Abstract

The expression of microRNAs (miRNAs) is dysregulated in many types of cancers including osteosarcoma (OS) due to genetic and epigenetic alterations. Among these, *miR-34c*, an effector of tumor suppressor P53 and an upstream negative regulator of Notch signaling in osteoblast differentiation, is dysregulated in OS. Here, we demonstrated a tumor suppressive role of *miR-34c* in OS progression using *in vitro* assays and *in vivo* genetic mouse models. We found that miR-34c inhibits the proliferation and the invasion of metastatic OS cells, which resulted in reduction of the tumor burden and increased overall survival in an orthotopic xenograft model. Moreover, the osteoblast specific over expression of *miR-34c* increased survival in the osteoblast specific p53 mutant OS mouse model. We found that *miR-34c* regulates the transcription of several genes in Notch signaling (*NOTCH1, JAG1* and *HEY2*) and in p53 mediated cell cycle and apoptosis (*CCNE2, E2F5, E2F2* and *HDAC1*). More interestingly, we found that the metastatic free survival probability was increased among a patient cohort from TARGET OS which has lower expression of direct targets of *miR-34c* that was identified in our transcriptome analysis such as *E2F5* and *NOTCH1*. In conclusion, we demonstrate that *miR-34c* is a tumor suppressive miRNA in OS progression *in vivo*. In addition, we highlight the therapeutic potential of targeting miR-34c in OS.

## INTRODUCTION

Osteosarcoma (OS) is the most common primary malignant bone cancer, comprising approximately 20% of all bone tumors and about 5% of pediatric tumors. It is the third most common type of cancer among children after leukemia and lymphoma. It has a bimodal distribution that peaks at under age 15 and after age 65 years (1,2). OS metastasizes to the lungs and has a high relapse rate. Five-year survival for patients with localized disease approaches 70%, but patients with pulmonary metastasis experience poor survival rate of 19-30% (3). The genetic causes of OS are extensive but high penetrance mutations have been found in familial cancer syndromes associated with *RB1, TP53, RECQL4* and *WRN* (4). In mice, the loss of p53 or Rb1 in osteoblastic lineages led to the spontaneous OS (5-7). Moreover, our group showed that gain of Notch1 in osteoblasts was also sufficient to initiate spontaneous OS (8). We and others have shown that Notch1 gain of function in early osteoblastic lineage cells leads to osteosclerosis due to proliferation of immature osteoblasts and inhibition of their differentiation into mature osteoblasts (9-12). Long term, this also led to the development of osteogenic sarcomas that mimicked human OS in terms of distribution and molecular signatures (8). As Notch signaling is also up-regulated in human OS samples, our transcriptional profiling of OS from p53 mutant mice also exhibited activation of Notch signaling (13).

Tumor suppressor p53 protein not only regulates the transcription of coding genes but also regulates the transcription of non-coding RNAs such as microRNAs (miRNAs). Among these, the evolutionally conserved miR-34 family (miR-34a, b, and c) has been reported as a tumor suppressor in response to DNA damage and oncogenic stress and induces apoptosis and cell-cycle arrest (14,15). Interestingly, downregulation of *miR-34* expression was seen in many types of human cancers such as colon, glioma, and non-small cell lung cancers (16-18). In addition, it has been shown that genetic and epigenetic alterations in OS lead to decreased *miR-34* expression level (19). These findings suggest an important role of miR-34 as a downstream factor of p53 and a potential tumor suppressor miRNA in OS, although this has not been tested *in vivo*.

In our previous study, we found that *miR-34c* was one of the significantly up-regulated miRNAs during BMP-2 mediated C2C12 osteoblast differentiation. More interestingly, we and others elucidated its essential role in bone homeostasis using *Col1a1 2*.*3kb-miR-34c* transgenic mouse model (20,21). We showed that the gain of miR-34c in osteoblasts regulates osteoclastogenesis via non-cell autonomous manner, which phenocopies the osteoblast-specific loss-of-function Notch1 mouse model (20). Mechanistically, we confirmed that miR-34c directly targets 3’ UTR of multiple components of Notch signaling (*Notch1, Notch2* and *Jag1*) to regulate bone homeostasis in the transgenic mouse model (20). We, therefore, hypothesized that the p53 downstream effector miR-34c targets Notch signaling and other pathways/targets to inhibit OS tumorigenesis.

In this study, we demonstrated the tumor suppressive role of miR-34c by using an orthotopic xenograft model and a genetic mouse model of the osteoblast-specific miR-34c gain-of-function on a spontaneous OS osteoblast-specific p53 mutant background. Mechanistically, we showed that miR-34c regulates Notch signaling and other cancer related pathways by targeting multiple genes (*CCNE2, E2F2, E2F5*, and *HDAC1*). Moreover, we found that the metastasis-free survival was increased among TARGET OS cohort, which had lower expression of *E2F5* and *NOTCH1*. Overall, our findings provide genetic support for a tumor suppressive function of miR-34c in mouse and human OS. In addition, our study highlights the therapeutic potential for targeting miR-34c in OS progression.

## RESULTS

### 1. miR-34c expression is reduced in the metastatic human osteosarcoma cell lines while Notch signaling components are activated in complementary fashion

The expression of miR-34 family is dysregulated in many cancers due to genetic alterations and epigenetic modifications. Therefore, we first accessed the expression level of *MIR-34c* in human osteosarcoma (OS) cell lines. *MIR-34c* expression was significantly reduced in 143B cells, a highly metastatic human OS cell line, compared to low-metastatic potential OS cell lines such as SJSA1, U2OS and MG63 (Figure 1A). Consistent with our finding, others have reported that *MIR-34c* expression in OS cell lines was decreased compared to human fetal osteoblast cells (hFOB) (22). In addition, we found Notch signaling components including *NOTCH1, NOTCH2* and *JAG1* expression were significantly elevated in high metastatic potential143B cells compared to low-metastatic potential human OS cell lines (Figure 1B, C and D). This expression pattern demonstrates an inverse correlation of miR-34c vs. Notch expression in the metastatic human OS cell line. This result is consistent with our previous report demonstrating miR-34c’s direct inhibition of Notch signaling during physiological osteoblast differentiation (20). Given that miR-34c is a downstream effector of p53 and an upstream negative regulator of Notch signaling, we hypothesized that miR-34c plays a tumor suppressive role in OS progression.

**Figure 1.**
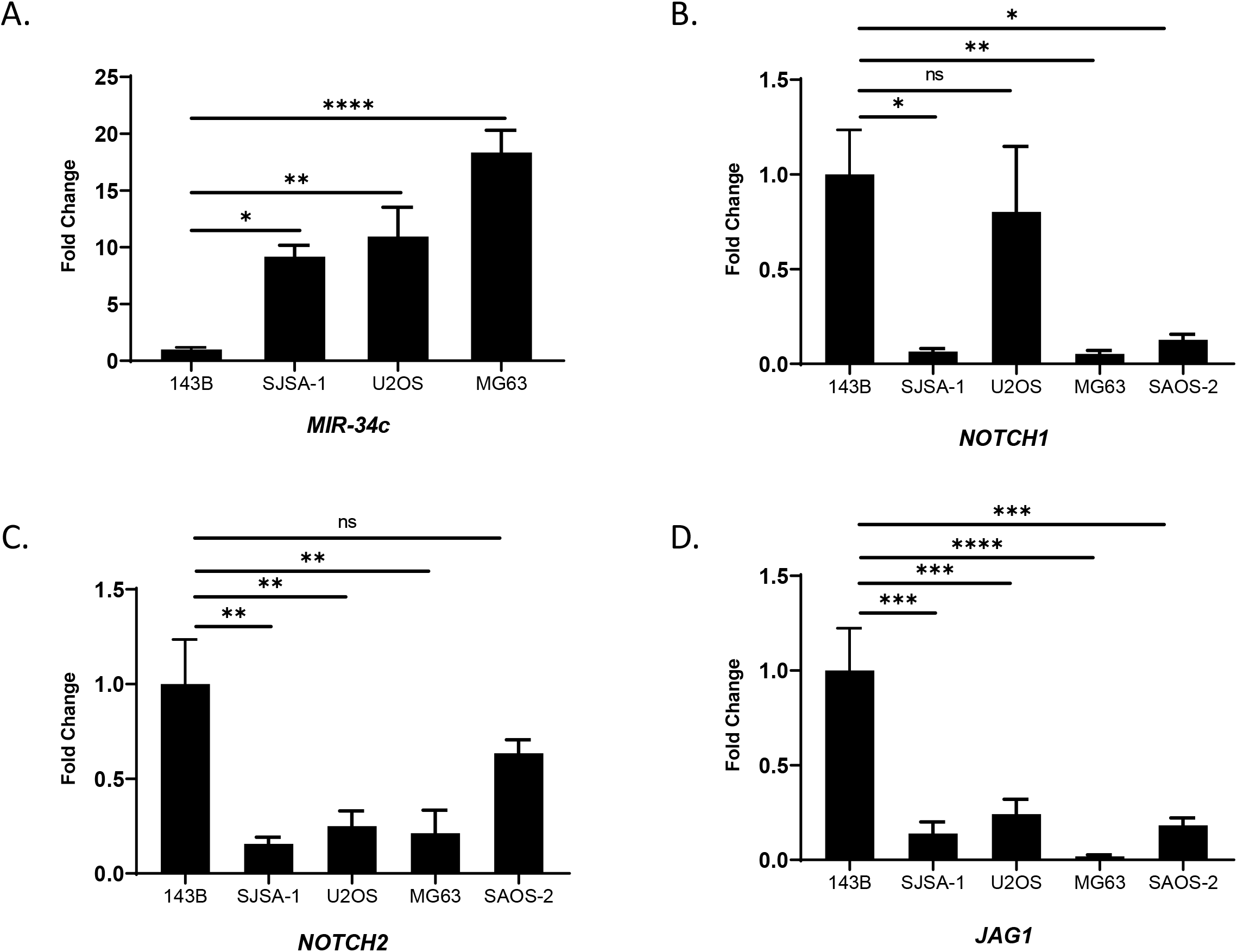
Expression of *MIR-34c* and Notch signaling components in human osteosarcoma (hOS) cell lines. qRT-PCR analysis of (A). *MIR-34c*, (B). *NOTCH1*, (C). *NOTCH2* and (D). *JAG1* expression in hOS cell line (143B, SJSA-1, U2OS, MG63 and SAOS-2). Values are Mean ± SEM, N=4. *, *P*<0.05; **, *P*<0.01; ***, *P*<0.001; ****, *P*<0.0001 (One-way ANOVA and Sidak test for multiple comparison with 143B)

### 2. miR-34c inhibits proliferation and invasiveness of metastatic human OS cell line

To test the tumor suppressive potential of *miR-34c* in OS, we generated a series of clonal cell lines of overexpressing *miR-34c* (LG34C: Luciferase-GFP-miR-34c) and their scramble control (LGS: Luciferase-GFP-Scramble) from 143B cells. The construct contains luciferase to enable live cell imaging, the quantitative analysis of tumor growth, and lung metastasis over time using bioluminescence imaging (Figure S1A). After establishing eight clonal cell lines from each LG34C and LGS control, we compared the luciferase activity by % luminescence (Figure S1B) and further confirmed the overexpression of *miR-34c* and the inverse expression of *NOTCH1* in LG34C lines vs. LGS control lines (Figure 2A and B). The cellular effects of the gain-of-miR-34c in 143B cells were then assessed. Overexpression of *miR-34c* (LG34C) resulted in decreased cell proliferation at 48 hours (hrs) (Figure 2C) and reduced invasion at 24 and 48 hrs compared to the LGS (Figure 2D). These cellular phenotypes suggest that *miR-34c* can affect the tumor burden and the metastatic property in OS progression.

**Figure 2.**
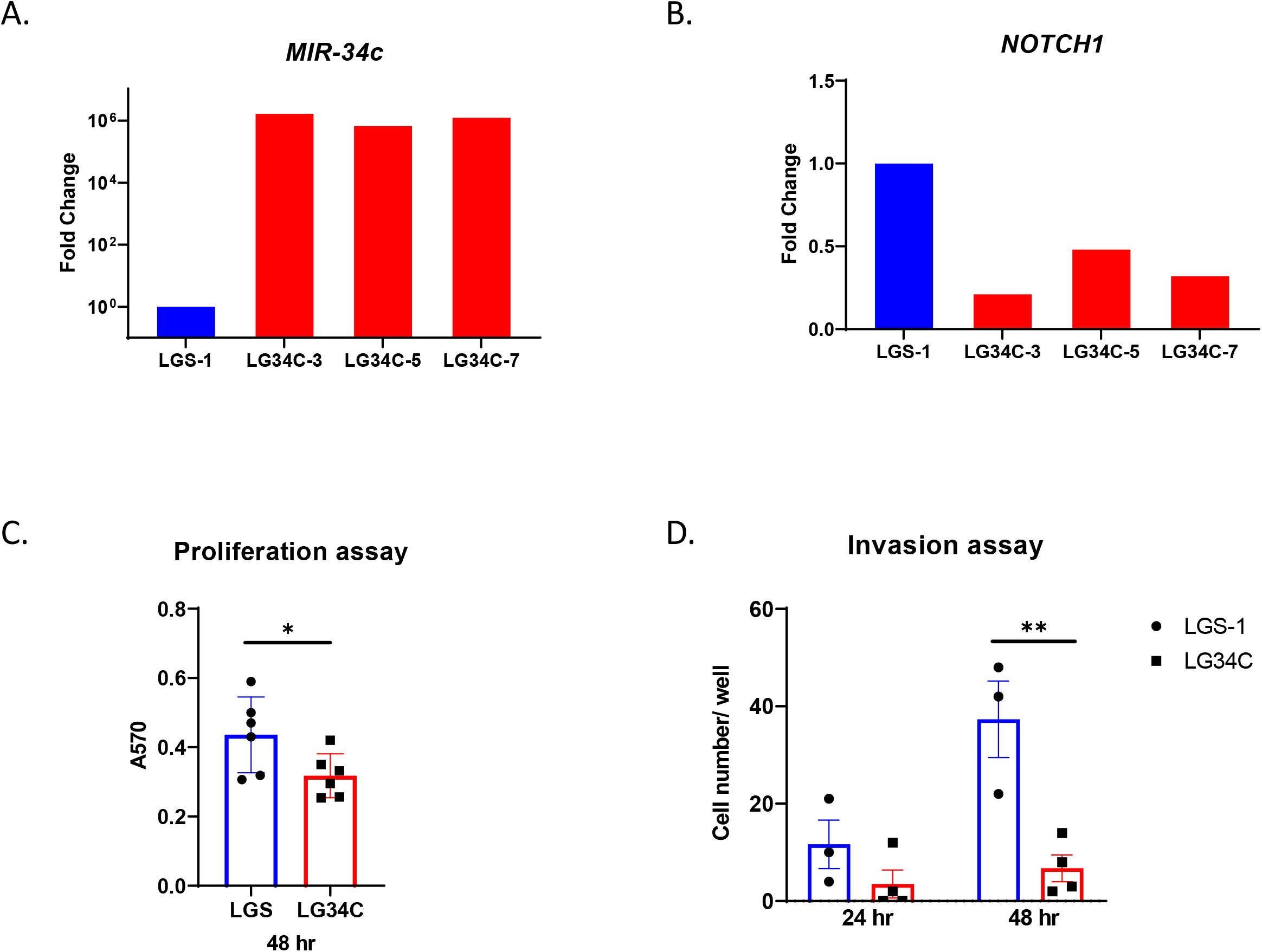
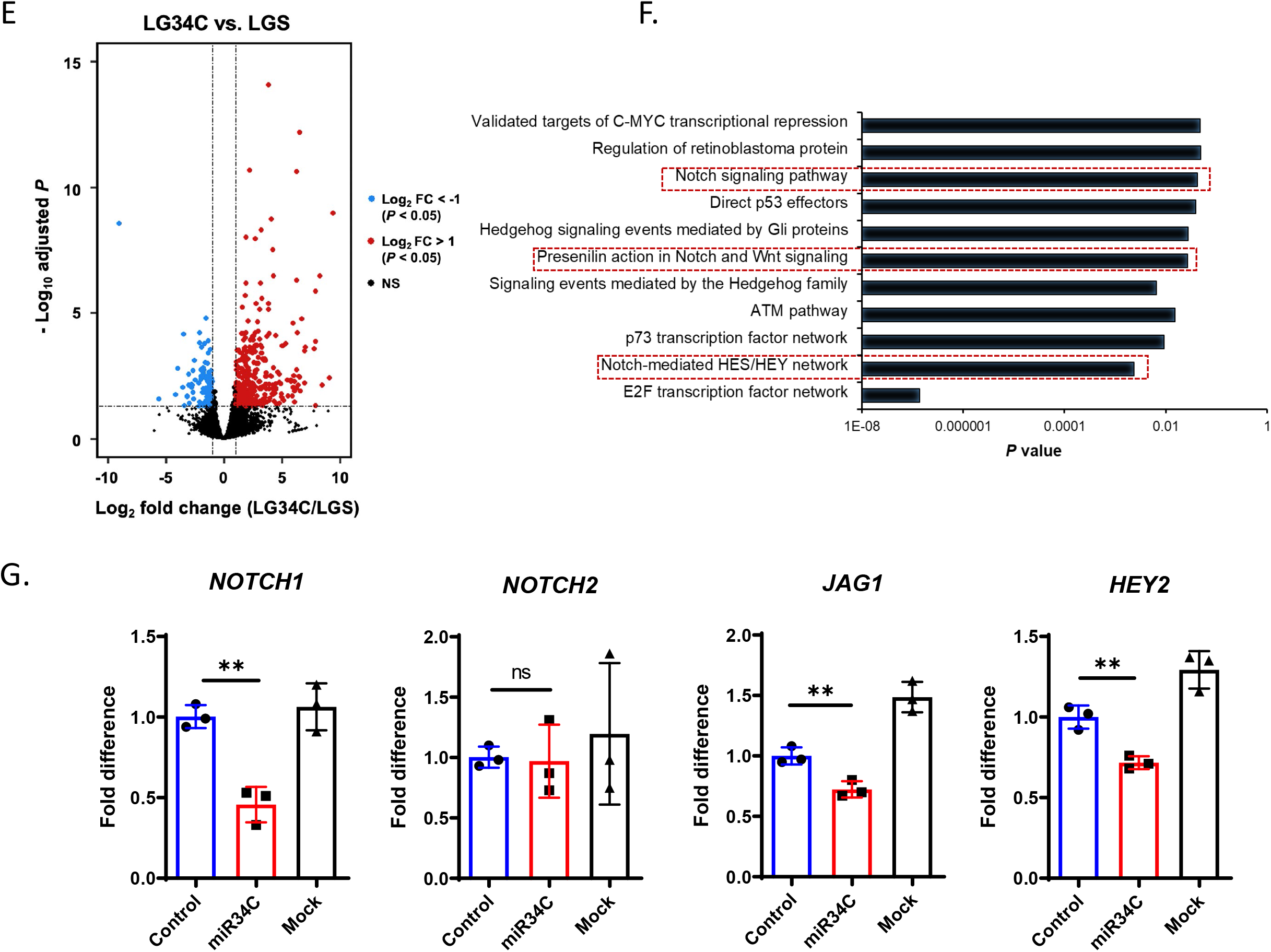
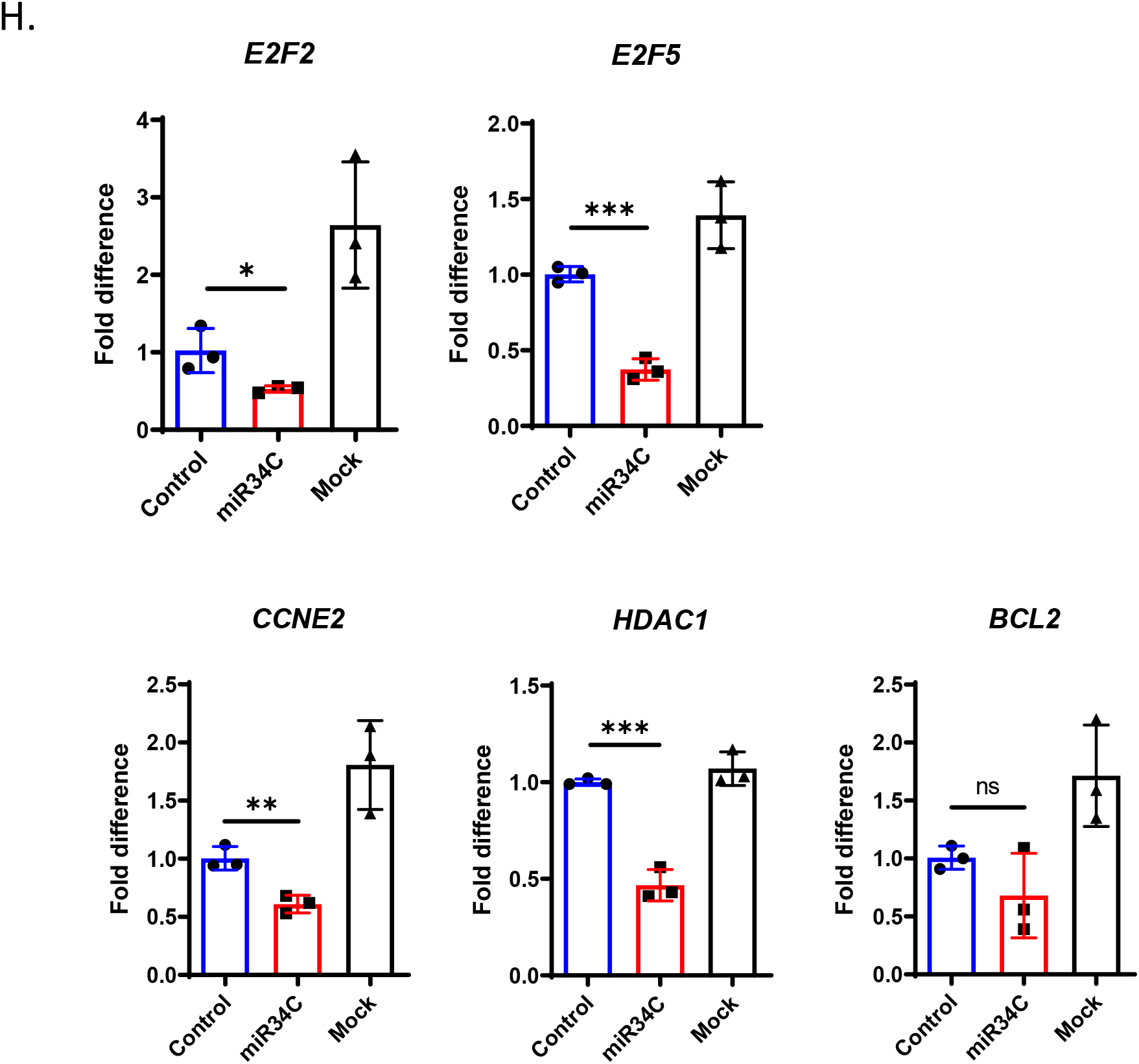
Gain-of-function *MIR-34c* in 143B decreases cell proliferation and invasion. (A). *MIR-34c* and (B). *NOTCH1* expression in each clonal stable 143B by qRT-PCR (n=1). (C). Decreased cell proliferation in *MIR-34c* expressing 143B cells by MTT assay at 48 hrs. Values are Mean ± SD, N=6. *, *P*<0.05 (Student t-test). (D). Decreased invasion in *MIR-34c* expressing cells at 24 and 48 hrs. Values are Mean ± SD, N=3. **, *P*<0.01(Student t-test). LGS (Luciferase-GFP-Scramble control stably expressing in 143B) and LG34C (Luciferase-GFP-miR-34c stably expressing in 143B). (E). Volcano plot of differentially expressed genes between LG34C and LGS with a Log_2_ fold change +/-1 and *P* value cutoff 0.05. Red dot indicates up-regulated genes and blue dot indicates down-regulated genes. FC (fold change). (F). Pathway affected in LG34C vs LGS within downregulated genes. (G). qRT-PCR analysis of Notch signaling pathway genes and (H). E2F transcription networks genes from 143B cells transfected with miR-34c mimic (miR34C) or control mimic (Control). Values are Mean ± SD, N=3.*, *P*<0.05; **, *P*<0.01; ***, *P*<0.001 (Student t-test comparing control vs miR34C group).

### 3. miR-34c regulates Notch signaling including cancer related pathways in OS tumorigenesis

To elucidate the underlying molecular mechanisms of tumor suppressive role of miR-34c in OS progression, we performed 3’-mRNA seq using three biological replicates of each clonal cell line (Control group: LGS1, LGS4 and LGS8; miR-34c group: LG34C3, LG34C5 and LG34C7). We obtained significantly altered expression of 110 downregulated genes (*p*<0.05 and Log_2_ fold change < -1) and 418 upregulated genes (*p*<0.05 and Log_2_ fold change > 1) (Figure 2E). Subsequently, we focused on the differentially downregulated genes to identify putative direct targets of *miR-34c*. Among downregulated genes, we found 22 genes were candidate direct targets of *miR-34c* based on the TargetScan analysis (Supplementary Table 1). Among affected pathways based on GO analysis of these downregulated genes, *E2F transcription factor network* seemed most significantly affected (*p* value = 1.38E-07) (Figure 2F). In addition, Notch signaling (*Notch-mediated HES/HEY network, Presenilin action in Notch and Wnt signaling and Notch signaling pathway*) was found as a recurrently affected pathway (Figure 2F). We identified and confirmed several specific targets representative of these pathways independently by overexpressing *miR-34c* mimic in 143B cells (Figure S2). *NOTCH1, JAG1* and *HEY2* transcript were significantly downregulated (Figure 2G). This is consistent with our previous transcriptome analysis performed with calvarial tissue from mice expressing *miR-34c* from the *Type I Collagen* promoter (*Col1a1 2*.*3kb-miR-34c*), in which many cancer-associated pathways including Notch signaling was dysregulated (20). We also found that *Direct p53 effectors, E2F transcription factor network*, and *Regulation of RB protein* were differentially regulated and genes associated in these pathway-*E2F2, E2F5, HDAC1* and *CCNE2* were significantly downregulated. These changes were further confirmed by qRT-PCR (Figure 2H). *BCL2*, a direct *miR-34c* target, also showed reduced expression that trended toward significance. Overall, the transcriptome analysis and target analysis in OS cell lines showed that *miR-34c* regulates multiple cancer related pathways and molecular regulators including Notch signaling pathway.

### 4. miR-34c overexpression reduces tumor burden *in vivo* in a xenograft model of OS

To determine the tumor suppressive function of miR-34c *in vivo*, we performed a xenograft study by intra-tibial (IT) injection of *miR-34c* overexpressing LG34C or LGS control cell lines in immune-deficient mice (Figure S1C). We monitored tumor growth and lung metastasis by detection of luciferase expression by bioluminescence imaging (Figure S1A). Consistent with a decreased proliferation observed with overexpression of *miR-34c* in LG34C cells (Figure 2C), we found reduced tumor burden in LG34C compared to LGS engrafted immune-deficient mice (Figure 3A and B). More interestingly, we found the prolonged survival in the gain-of-function miR-34c (LG34C) vs. control (LGS) groups (Figure 3C). Since we previously found a significant impact on the invasiveness of LG34C cells (Figure 2D), we also accessed the pulmonary metastasis *in vivo*, which is a primary cause of death of OS patients. However, there was no difference in the burden of lung metastasis based on the level of luminescence from the lung samples at the termination in this aggressive immune-deficient model (Figure S1D). Overall, these collective results revealed that miR-34c is a tumor suppressor in OS by inhibiting cell proliferation and invasiveness *in vitro* and decreasing tumor burden, thereby, increasing overall survival *in vivo*.

**Figure 3.**
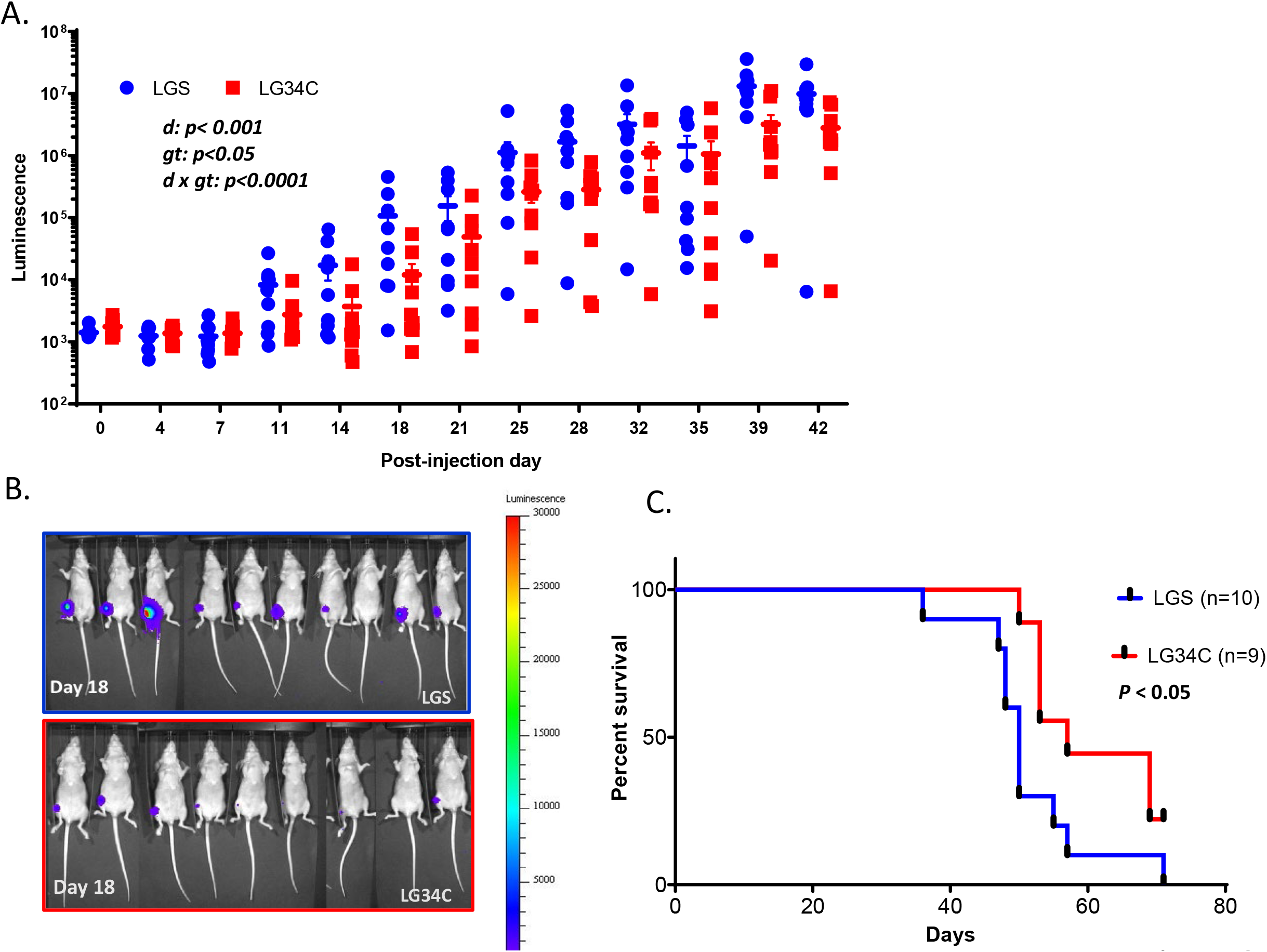
Gain of miR-34c significantly suppressed tumor growth *in vivo* and increased survival in xenograft model. (A). Tumor burden was quantified by luminescence intensity measured by *in vivo* whole animal bioluminescence imaging up to day 42 post-injection. N=9. *, *P*<0.05; ***, *P*<0.001; ****, *P*<0.0001 (Two-way ANOVA, d (Day), gt (Genotype)). (B). *In vivo* live bioluminescence imaging of tumors from each LGS and LG34C engrafted mice at day 18 post-injection. (C). Kaplan-Meier survival analysis showed increased percent survival of LG34C compared to LGS. *P*<0.05 (Log-rank test).

### 5. Gain of miR-34c in osteoblasts improves survival of a spontaneous genetic osteosarcoma mouse model (*Col1a1 rat 2*.*3kb Cre; p53* ^*f/f*^)

Deletion of TP53, or RB1 in the osteoblastic lineage cells led to the spontaneous osteosarcoma in the genetic mouse models (5-7). Moreover, human genetics show that the mutation or loss of these genes predispose to OS (4). Interestingly, the transcriptional analysis of OS from p53 mutant mice also showed significantly increased many components of Notch signaling pathway (13). In addition, we have demonstrated that the gain-of-function Notch1 in mature osteoblasts is sufficient to cause spontaneous OS in genetically engineered mouse model (8). More importantly, we found elevated Notch signaling in primary human OS samples and even higher in metastatic OS samples (13). To demonstrate the *in vivo* tumor suppressive role of miR-34c in OS, we over expressed *miR-34c* in the osteoblasts by intercrossing *Col1a1 2*.*3kb-miR-34c* transgenic mice with *p53* deletion in osteoblasts (*Col1a1 rat 2*.*3kb Cre; p53* ^*f/f*^: refer as *OB* ^*p53-/-*^) (7,20,23). We confirmed that *miR-34c* transcript level was elevated in the calvarial tissue of gain-of-function *miR-34c* transgenic mice in the absence of *p53* in the mature osteoblasts (*Col1a1 rat 2*.*3kb Cre; p53* ^*f/f*^; *Col1a1 2*.*3kb-miR-34c*: refer as *OB* ^*p53-/-; miR-34c*^) by qRT-PCR (Figure 4A). As expected from gain-of-function miR-34c, *Notch1* expression was decreased in the calvarial tissue of *OB* ^*p53-/-; miR-34c*^ (Figure 4A). We found that *OB* ^*p53-/-; miR-34c*^ mice showed increased survival rate compared to *OB* ^*p53-/-*^ indicating that *miR-34c* functions as a tumor suppressor in OS (Figure 4B). We also found a trend toward lower incidence of lung metastasis in *OB* ^*p53-/-; miR-34c*^ mice compared to *OB* ^*p53-/-*^ alone (Figure 4C). Overall, our genetic model of gain-of-function *miR-34c* in OS mouse model along with the orthotopic xenograft study confirms tumor suppressive miRNA in OS progression and suggests that miR-34c may affect lung metastasis.

**Figure 4.**
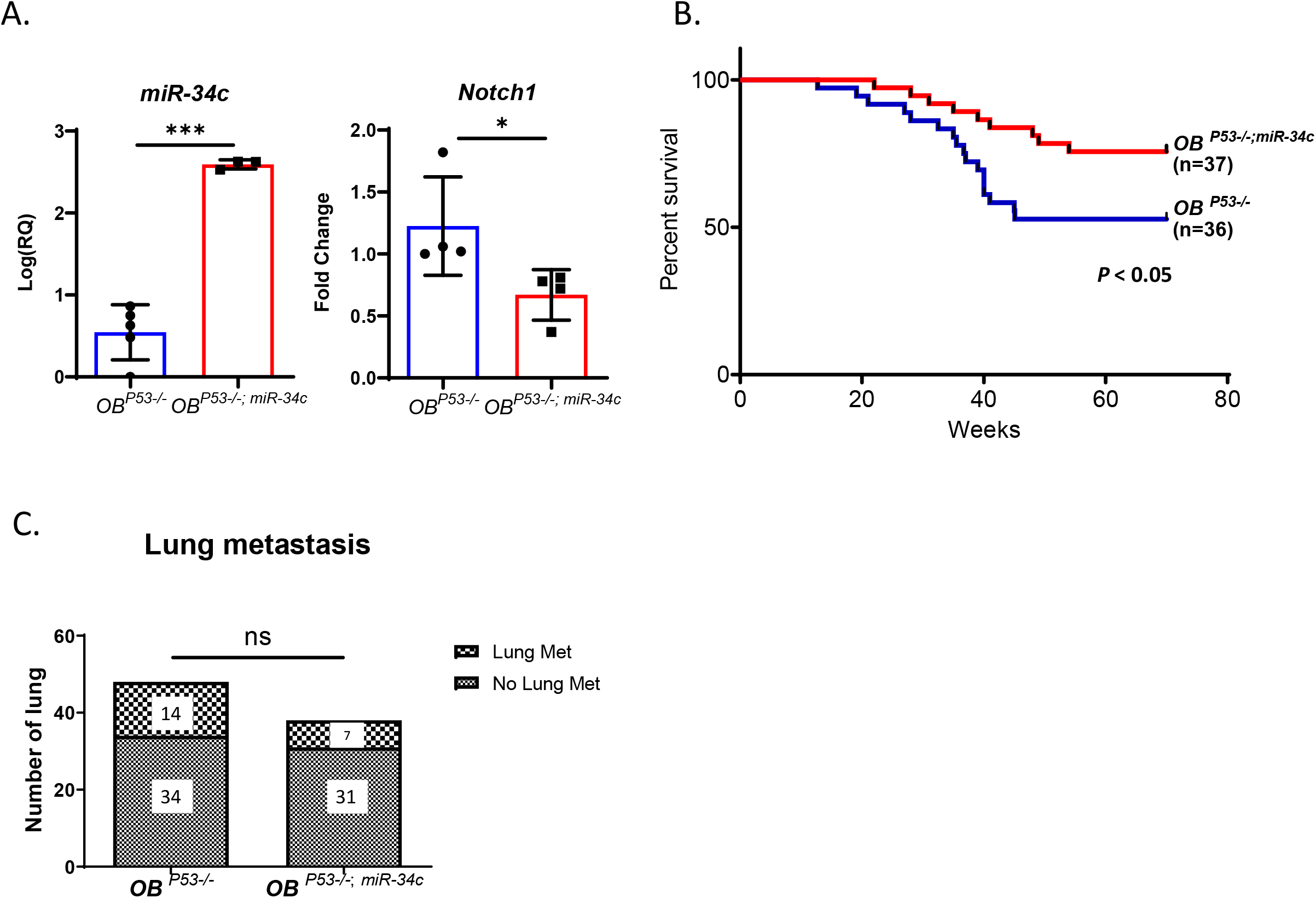
Genetic rescue by the osteoblast specific gain of *miR-34c* in *Col1a1 rat 2*.*3 kb Cre; p53* ^*f/f*^. (A). qRT-PCR analysis of *miR-34c* and *Notch1* from the calvarial tissue of each mouse line. Values are Mean ± SEM, N=5. *, *P*<0.05, ***, *P*<0.001 (Student t-test). (B). Kaplan-Meier survival analysis showed increased percent survival of *OB* ^*p53-/-; miR-34c*^ compared to *OB* ^*p53-/-*^. *P*<0.05 (Log-rank test). (C). No significant difference was found in the incidence of lung metastasis. ns (no significance) (Fisher exact test).

### 6. Increased expression of miR-34c targets in human OS patients associated with poor metastasis-free survival

In order to assess the clinical relevance of targets identified from the transcriptome analysis of stably over-expressing *miR-34c* 143B cells (Figure S2), we employed the TARGET OS cohort (n=87) and investigated the correlation between the expression level of these targets and the metastasis-free survival of OS patients. Kaplan-Meier survival analysis for these target genes, binned as tertiles, shows the metastasis-free survival probability of TARGET OS patient in Figure 5. Interestingly, we found higher survival rate of OS patients who have lower expression of *E2F5* and *NOTCH1* (Figure 5A and 5B). This metastasis-free survival analysis corroborates our findings of extended survival in *Col1a1 rat 2*.*3kb Cre; p53* ^*f/f*^; *Col1a1 2*.*3kb-miR-34c* (Figure 4B) and the xenograft models of OS progression (Figure 3).

**Figure 5.**
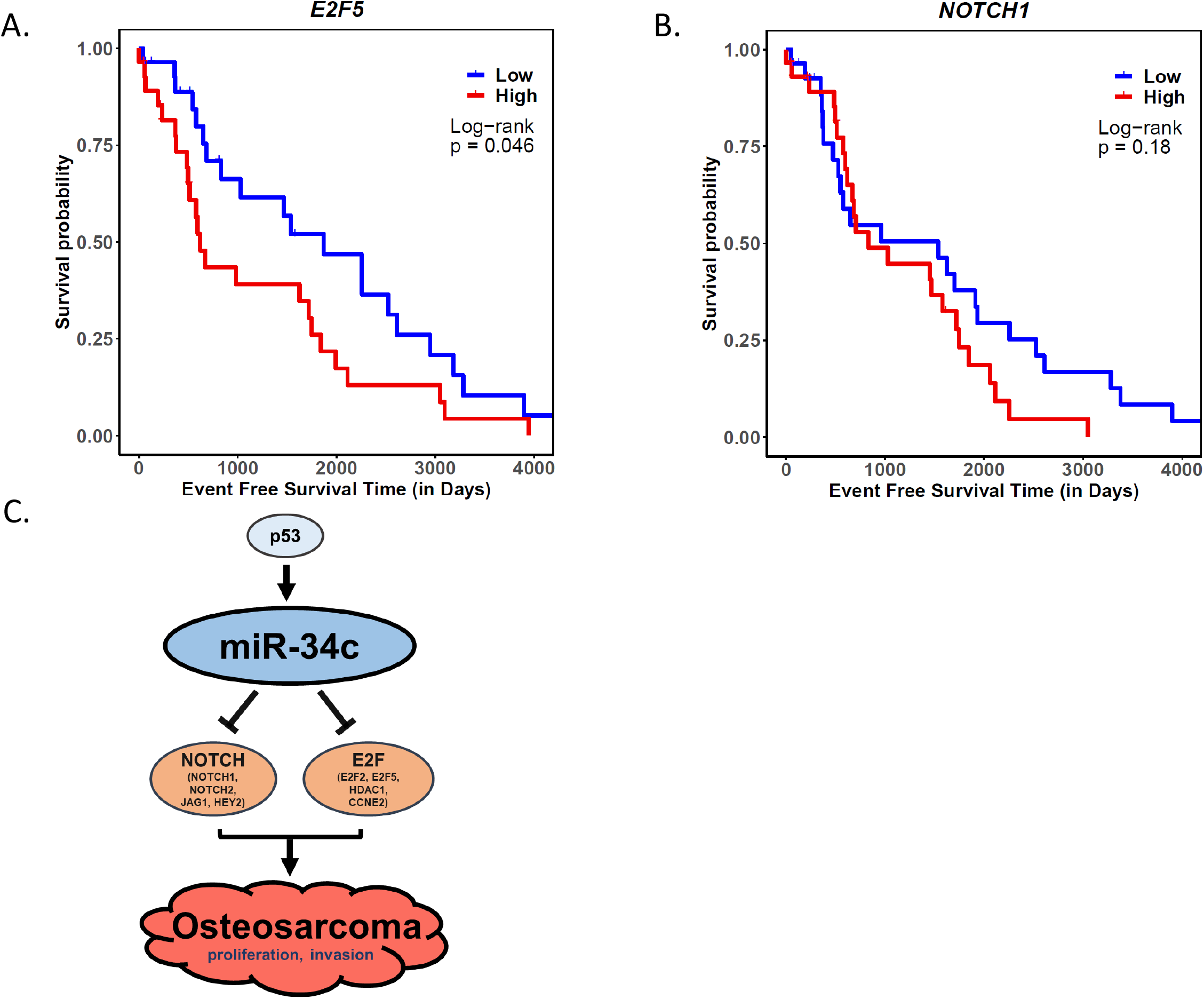
Analysis of miR-34c target expression and metastasis-free survival of the TARGET-OS patient cohort. (A). *E2F5* and (B). *NOTCH1* expression vs. metastasis-free survival of TARGET-OS patient cohort (N=87) by Kaplan-Meier survival analysis. (Log-rank test). (C). Schematic summary of tumor suppressive role of miR-34c by targeting Notch and E2F in OS progression.

In conclusion, our study demonstrated the tumor suppressive role of miR-34c in OS progression by targeting multiple pathways including Notch signaling and cell cycle regulators (Figure 5C). This was associated with decreased proliferative and invasive properties of OS cells *in vitro*. Moreover, we showed a tumor suppressive function by the genetic interaction of osteoblast specific gain-of-function miR-34c with p53 loss-of-function in a spontaneous OS mouse model. Finally, we identified clinical significance of miR-34c targets by correlating the OS patient survival with a subset of these targets using the TARGET OS cohort. Our findings provide genetic evidence in support of the tumor suppressive function of the miR-34c during OS progression and identify it as a potential therapeutic target for OS.

## DISCUSSION

Dysregulation of miRNAs by genetic and epigenetic alterations has been widely implicated in tumorigenesis, which contributes to their potential roles as oncogenes and tumor suppressors. Such alterations were also observed in OS compared to healthy bone (14). Among these, miRNA-34 family is subjected to genetic and epigenetic changes in OS. There is decreased expression of *miR-34c* due to gene deletions and to DNA hypermethylation (19). The restoration of miR-34 family in OS cell lines partially induced cell cycle arrest, apoptosis and delay proliferation and invasion, indicating a potential tumor suppressive role of miR-34 family. Interestingly, we also found that *miR-34c* expression was drastically suppressed in a highly metastatic human osteosarcoma cell line (143B) compared to low-metastatic potential cell lines (Figure 1) and the overexpression of *miR-34c* can suppress the proliferative and invasive properties of metastatic 143B cells. Previously, we and others demonstrated that miR-34c is one of the osteogenic miRNAs that also plays important roles in bone homeostasis (20,21). In this study, we further showed that miR-34c regulates Notch signaling by directly targeting many of its components (*Notch1, Notch2* and *Jag1*) in osteoblasts. Notch signaling is tightly regulated to maintain a balance of pools of progenitors vs. differentiated cells during bone homeostasis. Not surprisingly, dysregulation of Notch signaling has also been implicated in various cancers including OS. We found Notch signaling components were highly upregulated in primary and metastatic tumors from the OS patients (13). In addition, these components were highly elevated in OS from *p53* ^*+/-*^ mice suggesting the functional interaction of tumor suppressor P53 and Notch in OS progression. The definitive oncogenic role of Notch signaling in OS was further proven by spontaneous OS development in an osteoblast-specific gain-of-function Notch mouse model. Moreover, there was a synergistic effect between p53 and Notch in accelerating OS progression (8). Here, we assessed whether the physiological role of an upstream regulator during bone homeostasis may similarly serve as a tumor suppressor in the context of OS. *MiR-34c* appears to serve this dual role in bone homeostasis and in OS progression. Functionally, as miRNAs target many genes and pathways, *miR-34c* is a prime candidate for enhancing tumor suppressive function in OS.

Concepcion C. et al. showed that complete deletion of miR-34 family did not compromise normal development and the spontaneous tumorigenesis in mice (24). Furthermore, in contrast to p53-dificient mice, miR-34 family deficient animals did not display any increased susceptibility compared to irradiation-induced or c-Myc induced B cell lymphoma mice. Similarly, miR-34a alone had little or no effect on tumor formation in mice (25). It accelerated initiation and formation of tumors only when these mice were bred with other mice harboring oncogenic variant such as *KrasG12D* (Kras-driven lung cancer model) or *APC* (colon cancer mouse model) (25,26). Together, these loss-of-function mouse models suggest that miR-34 family is largely dispensable for p53 function and while not sufficient to initiate tumorigenesis, it may serve as an essential progression factor. To test the *in vivo* tumor suppressive role of miR-34c in OS pathogenesis, we crossed transgenic mice over expressing *miR-34c* in osteoblasts (*Col1a1 2*.*3kb-miR-34c*) to mice deficient for p53 in osteoblasts (*Col1a1 rat 2*.*3kb Cre; p53* ^*f/f*^ or p53 cKO). Gain of miR-34c in p53 cKO OS mice improved survival compared to p53 cKO OS mice, supporting the tumor suppressive role in OS progression (Figure 4B). Since the miR-34 family functions as a mediator of p53, our ectopic expression of *miR-34c* in osteoblasts could partially compensate the loss of p53 function in OS. Consistent with this finding, our xenograft study showed the decreased tumor burden and increased survival from the mice engrafted via intra-tibial injection with 143B cells stably expressing the miR-34c (LG34C) vs. scramble control (LGS) (Figure 3). However, we did not observed any improvement in pulmonary metastasis in either the genetic mouse model (Figure 4C) and xenograft studies (Figure S1D) despite significant decrease of invasiveness in miR-34c gain-of-function cells. The discrepancy may be explained partly by the difference of *in vivo* vs. *in vitro* sensitivity to *miR-34c* over expression and the complex tumor microenvironment involving extracellular matrix, blood vessels and host immune cells. Moreover, the aggressive nature of this orthotopic immune-deficient model may not provide sufficient time to progression to demonstrate a protective effect of *miR-34c* expression.

Mechanistically, Notch signaling and cancer related pathway were differentially regulated in *miR-34c* stably expressing 143B cells. This result is consistent with our previous study of the transgenic mouse model of overexpressing miR-34c in osteoblastic cells (*Col1a1 2*.*3kb-miR-34c*) (20). Many targets of miR-34c found in our transcriptome analysis from 143B cells were shared among different pathways. *E2F2* and *E2F5*, E2F family proteins, were significantly downregulated in *miR-34c* overexpressing 143B cells (Figure 2H). These factors directly bind to cell cycle associated factors and promote cell cycle progression (27). It is reported that *E2F5* was dysregulated in OS and shown to be downregulated by direct targeting of *miR-154-5p* in OS (28). *CCNE2*, involved in G0/G1 cell cycle regulation, was also downregulated in our transcriptome analysis, suggesting that inhibition of cell proliferation is one of the mechanisms of *miR-34c* mediated suppression in OS development. In addition, *HDAC1* was significantly downregulated in our transcriptome analysis. Treatment with panobinostat, a broad spectrum HDAC inhibitor, showed decreased OS growth and lung metastasis in an orthotopic xenograft model using human OS cell lines (29). In this report, treatment with the selective inhibitor for HDAC1/2 compromised the growth of OS *in vitro* and *in vivo*. Moreover, Notch signaling is critical in maintenance of cell proliferation, as suppression of Notch signaling by miR-34c, again, effectively targets the cellular proliferative phenotype.

Finally, we examined the potential for some of miR-34c targets to correlate with metastasis free survival of OS patient cohort from TARGET database. Notably, OS patients who have lower expression of *E2F5* among miR-34c targets show higher metastasis free survival (Figure 5A). Patients with lower expression of *NOTCH1* also trended to have a better prognosis (Figure 5B). Expression level of other miR-34c targets did not correlate with patient survival of TARGET OS cohort.

Collectively, our study demonstrated that miR-34c, is not only a critical miRNA for the maintenance of bone homeostasis, but also a tumor suppressor in OS progression. It can inhibit the proliferation and invasion of tumor cells *in vitro*. It also decreased tumor burden in an orthotopic model of OS. Finally, *miR-34c* overexpression in osteoblasts increased survival of the p53 cKO spontaneous OS model. Mechanistically, gain of miR-34c regulates many targets from cancer-related pathways and Notch pathway. Among these, *E2F5* may significantly correlated with human OS progression since its elevated expression is associated with poorer survival of a patient cohort from TARGET OS. Our findings highlight the significance of miR-34c targeting the multiple pathways/genes which could be candidates for OS therapy.

## MATERIALS AND METHODS

### Animals

*Col1a1 rat 2*.*3kb Cre; p53* ^*f/f*^ mice were kindly provided by Dr. Jason Yustein (Baylor College of Medicine, Houston, TX) (7). In brief, *Col1a1 rat 2*.*3kb Cre; p53* ^*f/f*^ is generated by crossing *Col1a1 2*.*3kb* rat promoter driving a *Cre* transgene in osteoblasts with *p53* ^*f/f*^ mice (23). To test the tumor suppressive role of *miR-34c* in OS, *Col1a1 2*.*3kb-miR-34c* transgenic mice (FVBN background) were crossed with *Col1a1 rat 2*.*3kb Cre; p53* ^*f/f*^ mice (20). All research performed with these mice was conducted in compliance with the Baylor Animal Protocol Committee (Baylor College of Medicine Animal Protocol AN5136). All animals had comprehensive necroscopies upon the time of euthanasia with complete dissection of tumor along with other organs including the lungs, liver and other bones. Portion of tumor and lungs were processed for the pathology for the histological analysis and lung metastasis.

### Cell lines and cell culture

The human osteosarcoma cell lines - 143B, SJSA1, SAOS2, U2OS and MG63 (CLS Cat# 300441) - were purchased from ATCC. 143B and MG63 cells were maintained in Minimum Essential medium (MEM) (GIBCO). SAOS2 and U2OS cells were maintained in McCoy’s 5A medium. SJSA1 cells were maintained in RPMI-1640 medium. All media were supplemented with 10% fetal bovine serum (FBS) (GIBCO) and 1% penicillin and streptomycin (GIBCO).

Stable cell lines of LG34C or LGS were established by sequentially infecting lentivirus expressing luciferase followed by lentivirus expressing either miR-34c-GFP or SCR-GFP in 143B parental cells. In brief, the pLenti PGK V5-LUC Neo (Addgene plasmid # 21471) was obtained from Addgene (30,31). The pGIPZ was purchased from Open Biosystems (vector # RHS4346). A 370 bp fragment of genomic DNA containing miRNA miR-34c was cloned into the *Xho I* and *Mlu I* restriction sites of the pLKO.1 vector (Addgene plasmid #52920) to express *miR-34c* (32). The non-silencing control of the pGIPZ vector served as the scramble control. Neomycin (125 mg/ml) and puromycin (1 μg/ml) were treated for the bulk selection. Further single clone from each cell line with a similar level of luciferase intensity was expanded to establish stable single clonal cell line of LG34C or LGS.

### Invasion assay

The invasion assays were conducted in 24-well plates with permeable supports on 8 μm Pore Polycarbonate Membrane (Corning Transwell 3422) for 24 and 48 hours. 8 μg/ul of Matrigel matrix (Sigma-Aldrich, E1270) was coated in the permeable supports (upper chambers) and incubated at 37°C for 1 hour. Cell suspensions prepared in serum free media were seeded onto the upper chamber (1×10^4^ cells/chamber) and culture media containing 10% FBS were loaded in the lower chamber. After the incubation, invaded cells in the lower chamber were washed with PBS and stained with 0.05% crystal violet for 1 hr. The invaded cells were captured under the microscope and counted in five random fields in triplicate.

### Cell proliferation assay

Cells were plated in 96-well plates (2500 cells/well) and grew for 48 hours. MTT reagent (MTT Cell Proliferation Assay, ATCC 30-1010K) were added into each well. Cells were then incubated with MTT reagent for 4 hours. After 4 hours of incubation, the media were removed and 100 ul of DMSO were added to resolve purple precipitate. After 30 minutes under the room temperature, the absorbance in each well was measured at 570 nm in a microtiter plate reader. Medium only wells were processed with the same procedure for the background reading. The average values of absorbance readings were subtracted with the average value of the blank.

### Gene expression analysis by qRT-PCR

Total RNA was extracted with TRIzol® (ThermoFisher Scientific), and cDNA was synthesized using Superscript III First-Strand RT-PCR kit (ThermoFisher Scientific). RNA expression was analyzed by qPCR with SYBR Green I reagent (Roche). *ACTB* was used as reference gene for the normalization. For miRNA qRT-PCR, TaqMan MicroRNA Assay (Applied Biosystems) were used to quantify the expression of mature miR-34c (Assay ID:000428). TaqMan Universal PCR Master Mix was used for amplification and detection. Sno202 RNA was used as reference gene for the normalization.

### Overexpression of miR-34c

143B cells were plated into a 6-well plate and transfected with 100 nM of miR-34c mimic or negative miRIDIAN mimic using Lipofectamine 2000 (Thermo Fisher Scientific) according to the manufacturer’s instructions. Cells were harvested 48 hours after transfection for targets’ mRNA analysis.

### RNA-Seq and data analysis

Total RNA was extracted from 143B stable cell lines (LGS and LG34C) using TRIzol® (ThermoFisher Scientific). Extracted RNA was then further purified by lithium chloride (ThermoFisher Scientific) precipitation. RNA quality and quantity were measured by Bioanalyzer (Agilent Technologies). For RNA-Seq, 250ng of total RNA was used for QuantSeq 3’ mRNA-seq library preparation (Lexogen) according to manufacturer’s instructions. The alignment was performed using HISAT2 (http://ccb.jhu.edu/software/hisat2/index.shtml) through Genialis (https://www.genialis.com) with hg19-ERCC as reference. Normalization, differential expression, hierarchical clustering, and Gene Ontology analysis were then performed using EnrichR (http://amp.pharm.mssm.edu/Enrichr/) provided from Genealis pipeline.

### Human data analysis using TARGET OS

The gene expression profiles and clinical information data of the OS patients were obtained from the Therapeutically Applicable Research to Generate Effective Treatments (https://ocg.cancer.gov/programs/target) initiative, phs000468. The RNA seq data of 87 osteosarcoma patient samples were obtained from TARGET data (https://portal.gdc.cancer.gov/projects). The z-scores of the gene expression (transcripts per million (TPM)) of TARGET OS patients were binned into the tertiles. Survival analysis of the top and bottom tertiles and metastasis-free survival time was performed with the Kaplan–Meier method using function survfit from package survival of R.

### Xenograft study

143B-LG34C or 143B-LGS cells were harvested and resuspended in cold PBS. For each injection, 10,000 cells were mixed with 50 ul Matrigel and intratibially injected into nude mice (10 mice / group). Tumor growth were monitored by the intraperitoneal (i.p.) injection of D-luciferin by *in vivo* whole animal bioluminescence imaging (IVIS Xenogen). Animals were imaged weekly to monitor tumor growth and lung metastasis.

### Statistical Analysis

To determine the statistical significance among groups, student t-test or analysis of variance (ANOVA) test were performed followed by multiple comparison test. \**P* < 0.05; \*\**P* < 0.01; \*\*\**P* < 0.005; \*\*\*\**P* < 0.0001; NS, not significant. Data are presented as mean ± SD (standard deviation) or mean ± SEM (standard error of mean). Kaplan-Meier calculations were used to show the overall survival time and the results were compared by a Log-rank (Mantel-Cox) test. Fisher’s exact test was used for the prevalence of lung metastasis in the genetic mouse model.

## ACKNOWLEDGMENTS

We would like to thank Dr. Jason Yustein for the initial discussion and sharing the valuable mouse line of *Col1a1 rat 2*.*3kb Cre; p53* ^*f/f*^ for our study; Lazar Kurenbekova for the initial help and teaching the intra-tibial injection to the nude mouse. This work was supported by the BCM Intellectual and Developmental Disabilities Research Center (P30 HD024064, U54 HD083092, and P50 HD103555) from the National Institute of Child Health & Human Development, CPRIT (Cancer Prevention and Research Institute of Texas) (RP170488) and the Rolanette and Berdon Lawrence Bone Disease Program of Texas.

## AUTHOR CONTRIBUTIONS

YB conceptualized the project, designed, performed the experiments and wrote the manuscript. HCZ and YTC assisted the xenograft study. HCZ supported transcriptome analysis and interpretation of data. SK performed TARGET OS analysis. EM and ZY assisted mouse colony maintenance and harvesting mouse. FG evaluated histological slides from primary tumor and lung. BL supervised the project.

