## Supplementary data for "*miRNA-34c* suppresses osteosarcoma progression *in vivo* by targeting Notch and E2F"

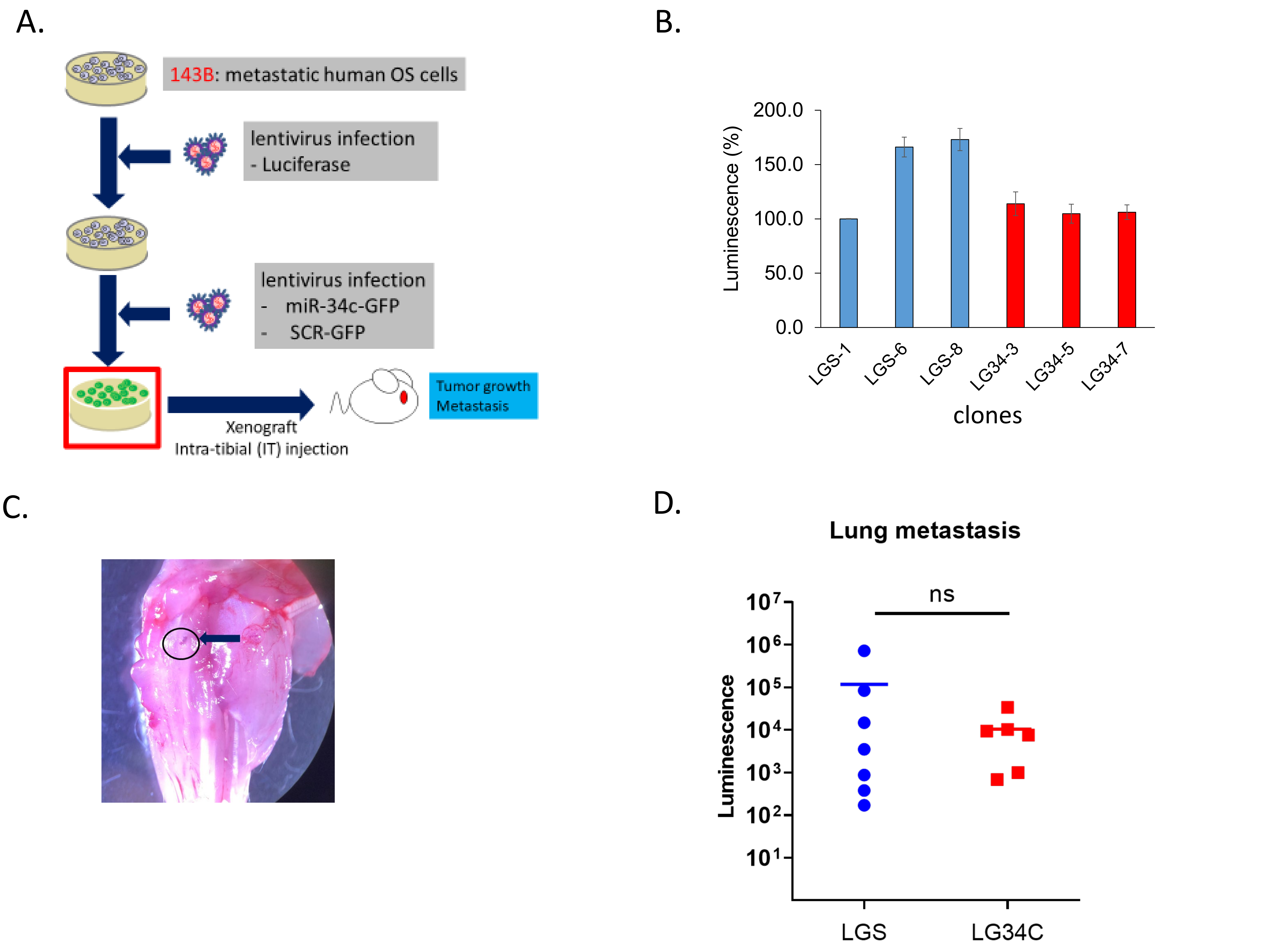
A.
143B: metastatic human OS cells
lentivirus infection
- Luciferase
lentivirus infection
- miR-34c-GFP
- SCR-GFP
Xenograft
Intra-tibial (IT) injection
Tumor growth
Metastasis
B.
Luminescence (%)
200.0
150.0
100.0
50.0
0.0
LGS-1
LGS-6
LGS-8
LG34-3
LG34-5
LG34-7
clones
C.
D.
Lung metastasis
ns
Luminescence
10^7
10^6
10^5
10^4
10^3
10^2
10^1
LGS
LG34C

Figure S1

| Term | Genes |
|---|---|
| E2F transcription factor network | **CCNE2;HDAC1**;MCM3;**E2F2;E2F5**;**CDC25A;TP73** |
| Notch-mediated HES/HEY network | **NOTCH1;HDAC1**;HEY2 |
| p73 transcription factor network | **CCNE2**;HEY2;**TP73** |
| ATM pathway | H2AFX;**CDC25A** |
| Signaling events mediated by the Hedgehog family | **GAS1**;GLI2 |
| Presenilin action in Notch and Wnt signaling | **NOTCH1;HDAC1** |
| Hedgehog signaling events mediated by Gli proteins | **HDAC1**;GLI2 |
| Direct p53 effectors | **BCL2;E2F2;TP73** |
| Notch signaling pathway | **NOTCH1;HDAC1** |
| Regulation of retinoblastoma protein | **HDAC1;E2F2** |
| Validated targets of C-MYC transcriptional repression | **HDAC1;BCL2** |

Figure S2

### SUPPLEMENTARY FIGURE LEGEND

**Figure S1. Effect of miR-34c in tumor growth and lung metastasis *in vivo* using luciferase reporter in orthotopic xenograft model.** (A). Flow chart of generating lentivirus mediated stable expression of miR-34c (LG34C) and scramble control (LGS) in 143B cells. (B). Percent luminescence from each clonal stable cells of LGS and LG34C. (C). Intratibial injection was performed to deliver LGS or LG34C to immune incompetent mice. (D). Lung metastasis was monitored at the termination by *ex vivo*. No significant difference was found (Student t-test).

**Figure S2. Downregulated genes in affected pathways.** Bolded genes in each pathway are directly targeted by miR-34c based on the TargetScan analysis.
